# Integrative analysis in head and neck cancer reveals distinct role of miRNome and methylome as tumour epigenetic drivers

**DOI:** 10.1101/2024.01.11.575141

**Authors:** Katarina Mandić, Nina Milutin Gašperov, Ksenija Božinović, Emil Dediol, Jure Krasić, Nino Sinčić, Magdalena Grce, Ivan Sabol, Anja Barešić

## Abstract

**Background:** Head and neck cancer is the sixth most common malignancy worldwide, with the relatively low 5-year survival rate, mainly because it is diagnosed at a late stage. Infection with HPV is well known etiology, which affects nature of these cancers and patients’ survival. Besides, it is considered that the main driving force for this type of cancer could be epigenetics. In this study we aimed to find potential epigenetic biomarkers, by integrating miRNome, methylome, and transcriptome analyses.

**Methods:** From the fresh head and neck cancer tissue samples, we chose a group for miRNome, methylome and transcriptome profiling, in comparison to adequate control samples. Bioinformatics analyses are performed in R v4.2.2. Count normalization and group differential expression for mRNA and the previously obtained miRNA count data was performed with DESeq2 v1.36. Gene set enrichment analysis was performed and visualized using gProfiler2 v0.2.1 Identification of miRNA targets was performed by querying in miRTarBase using multiMiR v1.18.0. Annotation of CpG sites merging into islands was obtained from RnBeads.hg19 v1.28.0. package. For the integrative analysis we performed kmeans clustering using stats v4.2.2 package, using 8-12 clusters and nstart 100.

**Results:** We found that transcriptome analysis divides samples into cancers and controls clusters, with no relation to HPV status or cancer anatomical location. Differently expressed genes (n=2781) were predominantly associated with signaling pathways of tumour progression. We identified a cluster of genes under control of the transcription factor E2F, that are significantly underexpressed in cancer tissue, as well as T cell immunity genes and genes related to regulation of transcription. Among overexpressed genes in tumours we found those that belong to cell cycle regulation and vasculature. A small number of genes were found significantly differently expressed in HPV positive versus HPV negative tumours (for example NEFH, ZFR2, TAF7L, ZNF541, and TYMS).

**Conclusions:** In this comprehensive study on an overlapping set of samples where the integration of miRNome, methylome and transcriptome analysis were performed for head and neck cancer, we demonstrated that the majority of genes were associated exclusively with miRNome or methylome and, to a lesser extent, under control of both epigenetic mechanisms.

## Background

Head and neck cancer is the sixth most common malignancy worldwide, with head and neck squamous cell carcinoma (HNSCC) accounting for more than 90% of cases. Unfortunately, HNSCC often gets diagnosed in a late phase when it is challenging to treat the tumour, which contributes to poor 5-year survival rate, currently at 66% (Johnson et al. 2020). Even though this type of cancer arises from a single cell type, i.e. the squamous cell, HNSCC is surprisingly of very heterogeneous nature (Canning et al. 2019). Primary site of tumour origin contributes to tumour heterogeneity, with most common sites being oral cavity, oropharynx, larynx, and sinonasal tract (Yan et al. 2011). Besides tumour location, the infection with high-risk human papilloma virus (HPV) types, especially in oropharyngeal area predominately affects HNSCC nature and patients’ survival (Kobayashi et al. 2018). HPV positive and HPV negative HNSCC can substantially differ in terms of etiology, genetics and epigenetics (Joseph and D’Souza 2012; Johnson et al. 2020). HPV-positive HNSCC mostly affect younger population, with no or low levels of smoking and/or alcohol use (Gillison et al. 2015).

Moreover, unlike the HPV-negative group, preferable site of tumour origin for HPV-positive HNSCC is oropharynx and it has been reported in many Western countries that this group of patients is associated with better overall survival and better response to therapy (Marur et al. 2010). There has been significant shift in the incidence of HPV-positive oropharyngeal squamous cancer in Western countries, where HPV is associated to HNSCC in almost 70% of cases (Marur and Forastiere 2016), and in fact, increase in 5-year survival of overall HNSCC cases could be contributed to higher prevalence of HPV-associated HNSCC (Johnson et al. 2020).

Up to date, many studies reported better outcomes for HPV-positive HNSCC (Perri et al. 2020; Kofler et al. 2014; Sun et al. 2021), however the consensus on specific biomarkers, which could improve diagnostic, prognostic and/or therapeutic approaches is still missing. Lack of consensus biomarkers could partially be owed to differences in studied populations sometimes inadequately registered tumour site of origin and possibly to the lack of stratification by the HPV status. Therefore, we consider the cohort presented below to be a specific case of tumours in terms of lifestyle, i.e. higher smoking and alcohol intake rates.

In this study, we aimed to identify epigenetic patterns in HNSCC since the main driving force for the HNSCC development is considered to be epigenetics, rather than genetics. We have previously analyzed the content and transcriptional levels of whole genome microRNA (miRNA) profile (miRNome) in HPV-positive and HPV-negative oral and oropharyngeal HNSCC samples using next generation sequencing and validated with qRT-PCR. We have also analyzed the whole genome methylation profile (methylome) in the same set of samples using the whole genome methylation array and validated by pyrosequencing. In this study, we assessed the whole genome transcriptome analysis on the same sample set and combined the mRNA, miRNA and methylation information to elucidate relevant epigenetic mechanisms and their interactions. Motivated by the limited literature regarding integrative studies on epigenetic changes, we set out to elucidate whether miRNAs or DNA methylation have more prominent impact on HNSCC development. Moreover, we aimed to identify key mechanisms and/or cellular signaling pathways of genes that have been differentially expressed in HNSCC compared to healthy controls. Consequently, we provide a better understanding of the epigenetic influence on the development and progression of HPV- positive and HPV-negative HNSCC, shedding light on mechanisms underlying prognostic, diagnostic and therapeutic strategies. In addition, integrative analysis points out key target genes and signaling pathways deregulated in HNSCC in Croatian and similar populations.

## Material and Methods

### Patient material

A total of 61 fresh oropharyngeal (OP) and oral (O) HNSCC primary tumour samples have been obtained from Dubrava Clinical Hospital in the 2015-2018 time period. Twenty-two of them and three non-malignant tonsil control samples were included in the miRNA profiling (Božinović et al. 2019), column 2 in **Table 1**. Of 22 sequenced samples, two were excluded, due to one sample being a recurrent tumour and one outlier. miRNome analysis and detailed patients’ socioepidemiologic and clinical data have been described in our previous study. From the previously published analysis of HNSCC methylome (Milutin Gašperov et al. 2020), of 22 samples included in miRNA study, 12 HNSCC samples were included in the whole genome methylation profiling (column 3, **Table 1**), as well as one additional sample that had methylome but no transcriptome and miRNome profile. Eight healthy swabs of oral mucosa were included in methylome profiling, which served as control samples.

**Table 1.**
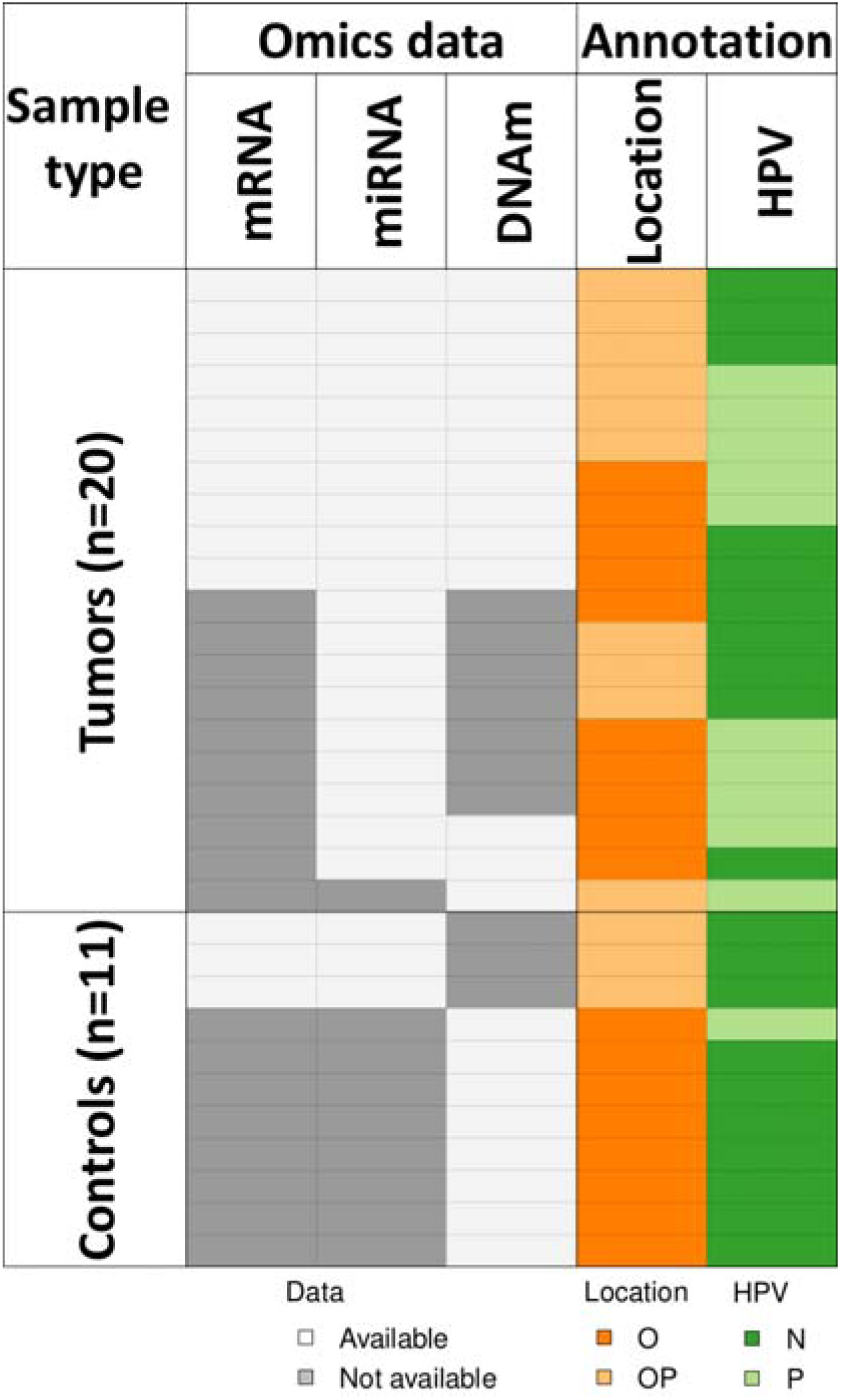
Samples used in integrative analysis. Sequenced samples are shown in white. O=oral, OP=oropharyngeal, P= HPV-positive, N=HPV-negative.

For the transcriptome profiling, we selected 10 fresh HNSCC samples (6 oral and 4 oropharyngeal) with previous miRNome and methylome data (column 1, **Table 1**). Three fresh tonsil tissues from the miRNome dataset were included in transcriptome profiling as control tissue. HPV status was assessed by PCR in all samples as described previously.

### Transcriptome profiling

As stated above, 13 samples were selected for transcriptome analysis in the current study to complement previously obtained omics results. Sample annotation for samples used in transcriptome analysis and subsequent integrative epigenetic study is presented in Table 1. Transcriptome was assessed using TrueSeq Stranded mRNA (Illumina) kit for library preparation, followed by sequencing on Nextseq500 instrument using NextSeq HIGH 150 cycles (Illumina) reagents on high output flow cell in paired end mode. Debarcoding was performed using Basespace (Illumina) platform. Quality control of sequencing data was performed with FastQC v0.11.9. Sequences were aligned to the human reference genome GRCh37 downloaded from GENCODE (released in 2013) and quantified using salmon v1.5.0, (Patro et al. 2017) reporting estimated fragment counts per transcript.

### Bioinformatics preprocessing

All bioinformatics analyses listed hereafter were performed in R v4.2.2, and refer to R language packages, unless otherwise stated. The preprocessing consisted of transcriptome gene count determination, reprocessing of miRNA counts (for a subset of samples from Božinović 2019) and reprocessing of differentially methylated CpGs (for a subset of samples from Milutin Gašperov 2020), here expanded to CpG islands.

Count normalization and group differential expression for mRNA and the previously obtained miRNA count data was performed with DESeq2 v1.36. Pre-filtering of the data set was performed to reduce genes that have nearly no information about gene expression by filtering out genes and miRNAs with total counts across all samples less than 10. Count variance stabilization transformation was performed with regularized logarithm transformation (DESeq2 rlog function) and used as input to calculate principal components with plotPCA function from DESeq2 and Euclidean sample distances with dist function from the stats v4.2.2 package. Counts normalization was performed with DESeq function with log fold change shrinkage. We calculated differential expression in tumour *versus* control and HPV positive *versus* negative samples (**Table 1**). Differentially expressed genes (DEGs) and miRNA (DEmiR) were selected with BH-adjusted p<0.05 set as significant.

Gene set enrichment analysis (GSEA) was performed and visualized using gProfiler2 v0.2.1 on DEGs using annotated human genes as the background set for the hypergeometric test.

We applied the tailor-made multiple test correction method Set Counts and Sizes (SCS) (Raudvere et al. 2019) and a p-value threshold of 0.05. GSEA was performed across multiple databases: Gene Ontology: Biological Processes (GO:BP) (Ashburner et al. 2000), Kyoto Encyclopedia of Genes and Genomes (KEGG) (Kanehisa et al. 2019), Reactome (Fabregat et al. 2018), miRTarBase (Chou et al. 2018) and TRANSFAC (Matys et al. 2006). We used these parameters for all subsequent GSEA.

Identification of miRNA targets was performed by querying significant DEGs against DEmiRs using multiMiR v1.18.0 and the miRTarBase database (Huang et al. 2022), selecting only experimentally validated miRNA-gene interactions.

Previously published DNA methylation (DNAm) data (Milutin Gašperov et al. 2020) was loaded with RnBeads v2.14.0. Differential methylation (DM) was calculated comparing tumour *versus* control groups on CpG sites and islands (CGIs). Annotation of CpG sites merging into islands was obtained from RnBeads.hg19 v1.28.0 package. All results are presented on CGIs unless otherwise stated. CGI DM was defined as the quotient of methylation between groups across all CpG sites in a CGI, with significance level set to FDR- adjusted p<0.05. Promoter region annotations were obtained from the RnBeads package, defined as 1500 bases upstream and 500 bases downstream of the transcription start site. CGI targets were assigned where CGI overlapped a promoter region.

### Integrative analysis

Workflow for comparing all omics data and identifying specific mechanisms in HNSCC is summarized in **Figure 1**.

**Figure 1.**
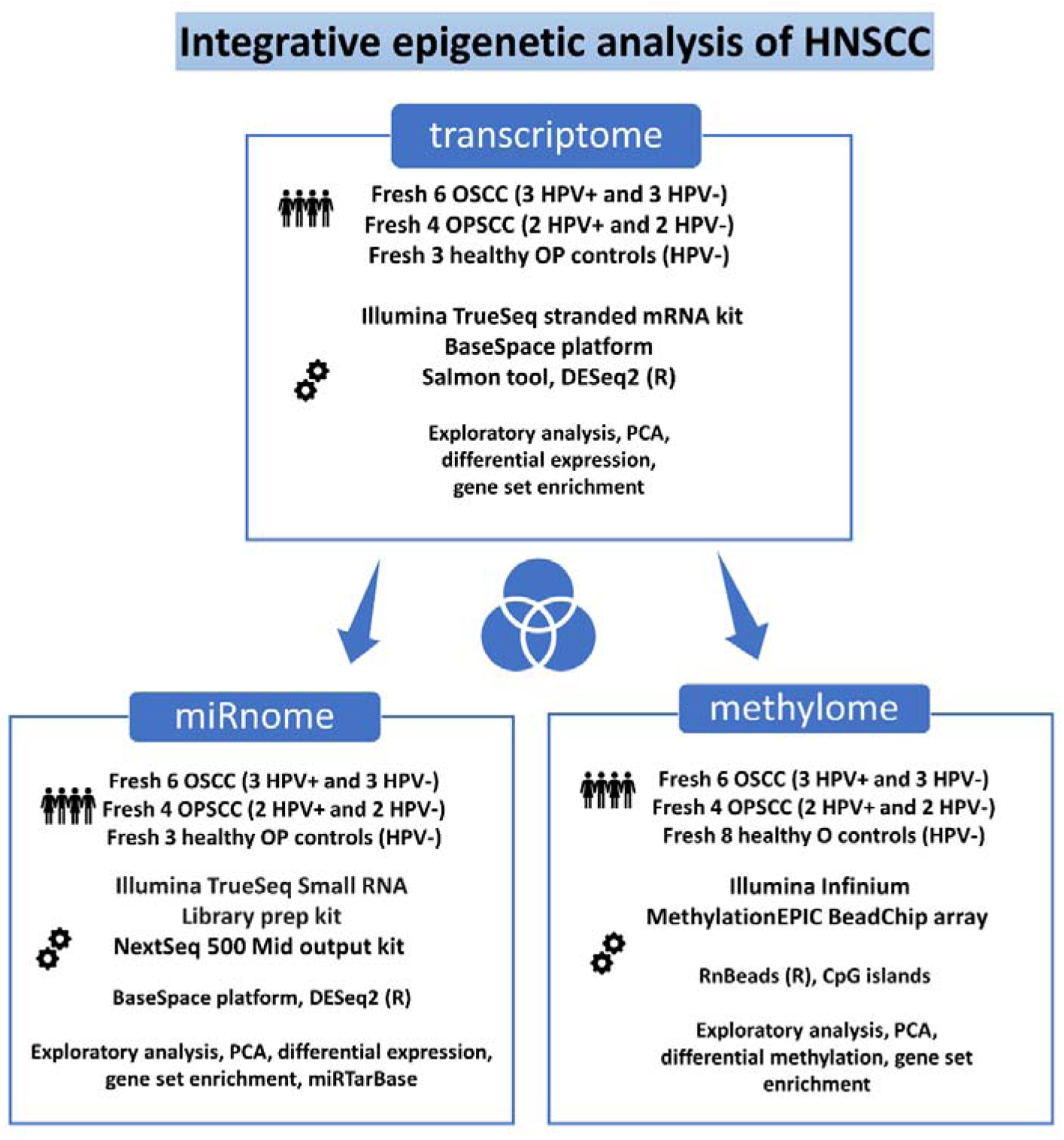
Schematic representation of integrative epigenetic HNSCC study.

Three -omics datasets were compared, centered on identified DEGs (**Supplemental table S1**). Multiple miRNA-gene pairs were reduced by selecting only the miRNA with highest absolute value of Spearman correlation of counts (mirRNA_Rho) for the given DEG. Multiple gene–CGI pairs were not as common, and the CGI with the lowest p-value was selected for every DEG. Resulting table contained DEGs and their associated miRNA and/or CGI. We assessed the interplay of DEmiRs, DM CGIs and DEGs in two ways: by obtaining simple overlap of DEGs targeted by DEmiRS or DM CGIs and kmeans clustering. The simple overlap was based on whether DEG’s expression levels were associated with DEmiR’s activity and/or CGI methylation levels. Kmeans clustering was performed on log-transformed, centered and scaled values of fold change for DEGs and DEmiRs, and DNAm quotient. We performed kmeans clustering using stats v4.2.2 package, using 8-12 clusters and nstart 100, with the best performing result for 11 clusters.

## Results

### Transcriptome profiling

The whole transcriptome profiling was performed on tissue samples from 13 individuals with different anatomical locations and HPV infection status (**Table 1**). The top two principal components of the transcriptome (**Figure 2a**), which explain 55% of the variance in the samples, splits samples into two clusters of tumour and control samples, with no obvious contribution of HPV status or tumour anatomical location. We observe the same pattern in miRNome and methylome data (**Figure 2b** and **c**).

**Figure 2.**
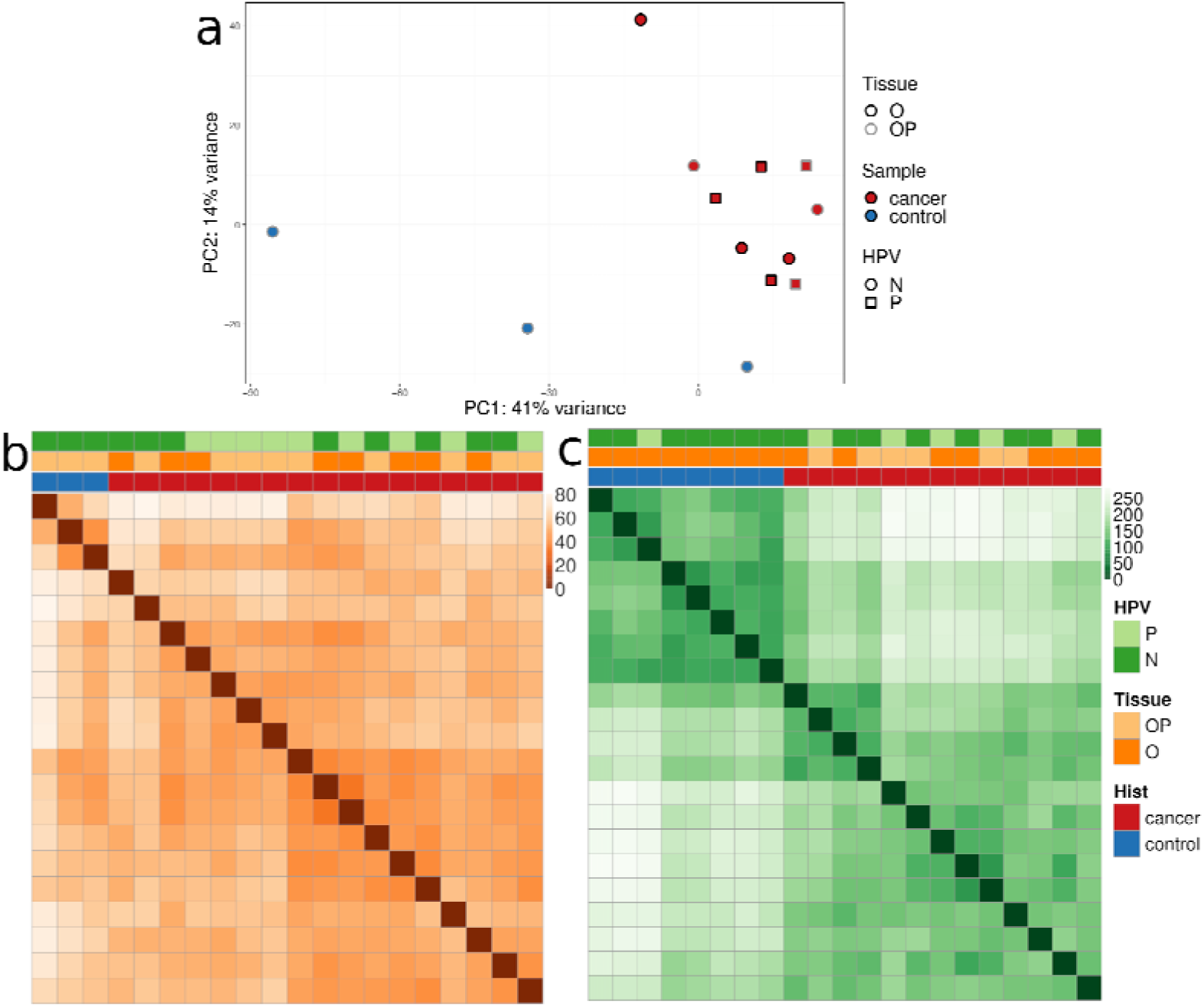
Exploratory analysis of three omics data sets: a) PCA plot for transcriptome data; b) sample distance heatmap for miRNome data; c) sample distance heatmap for methylome. N - HPV-negative, P - HPV-positive O - oral tumours, OP - oropharyngeal tumours.

### Differential expression and GSEA

Exactly 18104 genes had no less than 10 detected reads across samples. Differential expression analysis revealed that 1248 (6.9%) of genes were significantly overexpressed and 1533 (8.5%) were significantly underexpressed in tumour (BH-adjusted p<0.05), when compared to transcriptomes of control samples. GSEA on DEGs showed enrichment of signaling pathways associated with tumour progression, such as cell cycle regulation and oncogenic signaling pathways such as NF-kB (**Figure 3**), for a full list of terms see **Supplemental table S2**.

**Figure 3.**
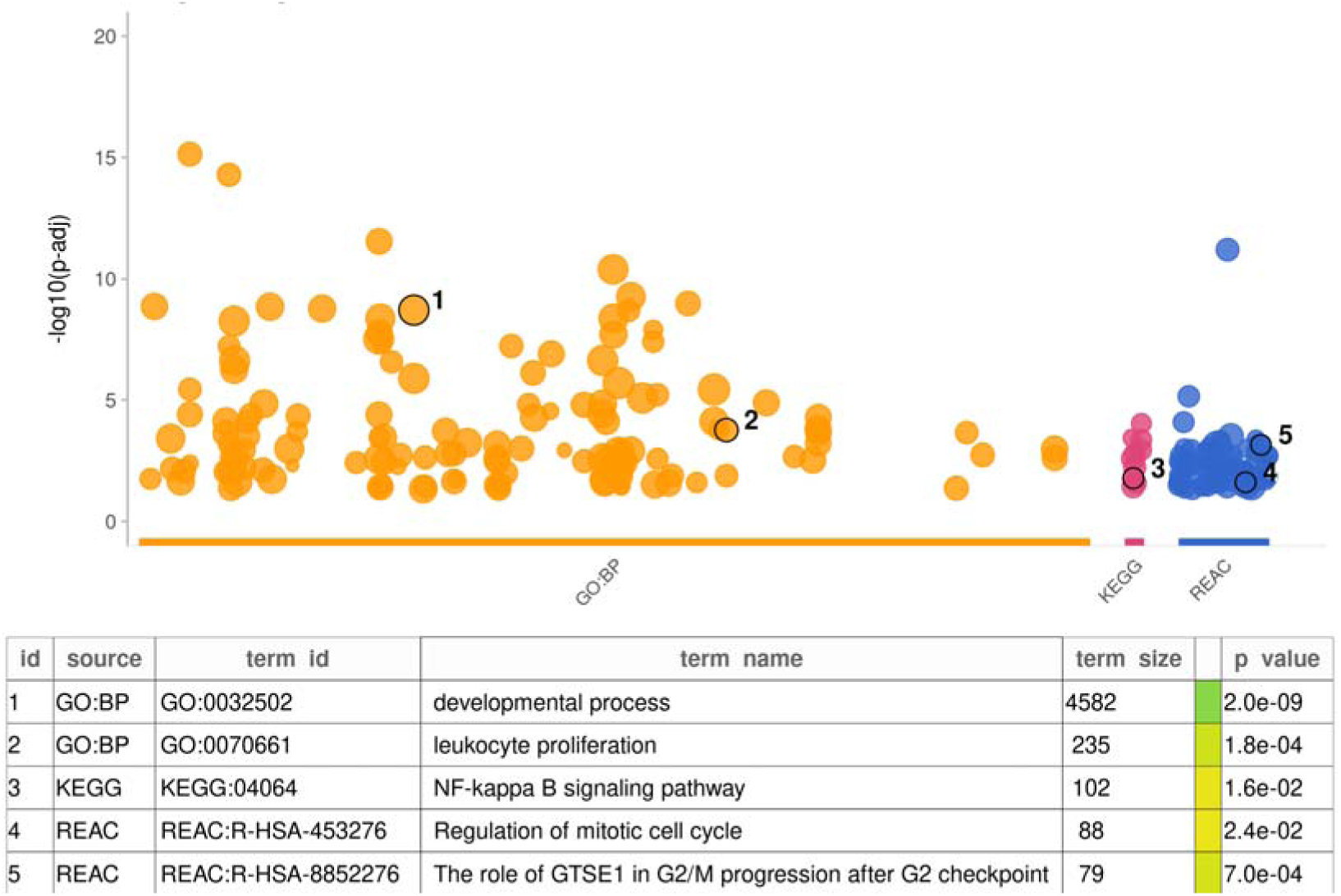
Gene enrichment analysis of tumour *vs.* control samples. Databases shown are GO:BP (orange), KEGG (pink), and REACTOME (REAC, blue). The highlighted terms correspond to key concepts associated with cancer progression and regulation.

### Differential miRNA and DNA methylation assessment

Integration of 514 significant DEmiRs based on MiRNome data (Božinović et al. 2019) with the database containing experimentally supported miRNA-target gene interactions yielded 3042 interactions among 90 different miRNAs and 1227 target genes, hereafter termed DEmiR-DEGs. In our analysis Most of these genes were regulated by a single miRNA, however some were associated with multiple miRNAs. Notably, two genes with the highest number of associated DEmiRs, *BCL2* and *MYC*, which play a role in regulating cell proliferation and apoptosis, and were associated with 18 and 16 miRNAs, respectively. In addition, some miRNAs were associated with many target genes, harboring potential therapeutic value by potentially affecting many cancer-related signaling pathways. An extreme example is a frequently reported tumour suppressor hsa-miR-26b-5p putatively forming interactions with 255 unique DEGs. Within our dataset hsa-miR-26b-5p showed a high negative expression correlation with known oncogenes such as *MMP10* (Rho= -0.81, p<0.005), *IGSF3* (Rho= -0.79, p<0.005) and *ARNTL2* (Rho= -0.79, p<0.005) (**Supplemental table S1**).

Overall methylation levels in our samples showed that more than 300,000 CpG sites were hypomethylated in tumour samples compared to controls (FDR-adjusted p<0.05), while 72,000 were hypermethylated (**Supplemental table S3**). We also noted a larger number of hypomethylated CGIs (6557) than hypermethylated (5457), although the difference is not as pronounced as in CpG sites (**Supplemental table S4**). The DM CpGs predominantly fell into gene bodies, but also some fell in intergenic regions corresponding to repeated elements. These regions consistently showed a larger number of hypomethylated CpGs compared to hypermethylated ones (**Supplemental figure S1**).

To identify deregulated tumour suppressors in our samples, we used the TSGene database 2.0 (Zhao et al. 2016) containing tumour suppressors curated from several thousand publications combined with expression and mutational profiles. We pinpointed 10 genes that were underexpressed and contained hypermethylated CGIs in the promoter region in our samples (**Supplemental table S5**).

To assess the effects of transcription factors on DEGs that had differentially methylated CGIs (DM-DEGS) within our dataset we performed GSEA using transcription factor database TRANSFAC. We found associations with several groups of transcription factors including E2F, SP/KLF and AP-2 families (**Supplemental table S6**).

Of note we found the enrichment of genes regulated by the E2F family of TFs (SCS-adjusted p<10), which play a critical role in cell cycle control. Furthermore, GSEA on the same dataset using the Reactome database showed enrichment of genes involved in cell cycle regulation such as G1/S Transition and G2/M Checkpoints (SCS-adjusted p<10^-4^), supporting the above findings.

### Integrative analysis: Influence of miRNome and methylome on transcription on transcriptional activity in HNSCC

Each DEG was grouped based on the association with DEmiR or DM CGI, shown in **Figure 4**. We obtained four groups: DEGs associated only with DEmiR (DEmiR-DEGs), DEGs associated only with DM CGIs (DM-DEGs), DEGs associated with both DEmiR and DM (DEmiR+DM- DEGs), DEGs not associated with either DEmiR or DM CGIs (noReg-DEGs). The groups had different proportion of differentially represented GSEA terms, i.e. GO:BP, KEGG, miRTarBase (MIRNA), Reactome and TRANSFAC (TF) (**Figure 4**, **Supplemental table S7**). The DM-DEGs group were enriched by genes regulated by TFs but no significant results of GO biological pathways was detected. The DEmiR-DEGs group were enriched with pathways related to signaling and immunological systems. The noReg-DEGs group was enriched with immunological processes and muscle system processes. DEmiR+DM-DEGs were enriched with general regulatory processes: biological, metabolic and cellular processes. These terms are high level terms in GO biological processes, suggesting that genes in the intersection have broad regulatory roles.

**Figure 4.**
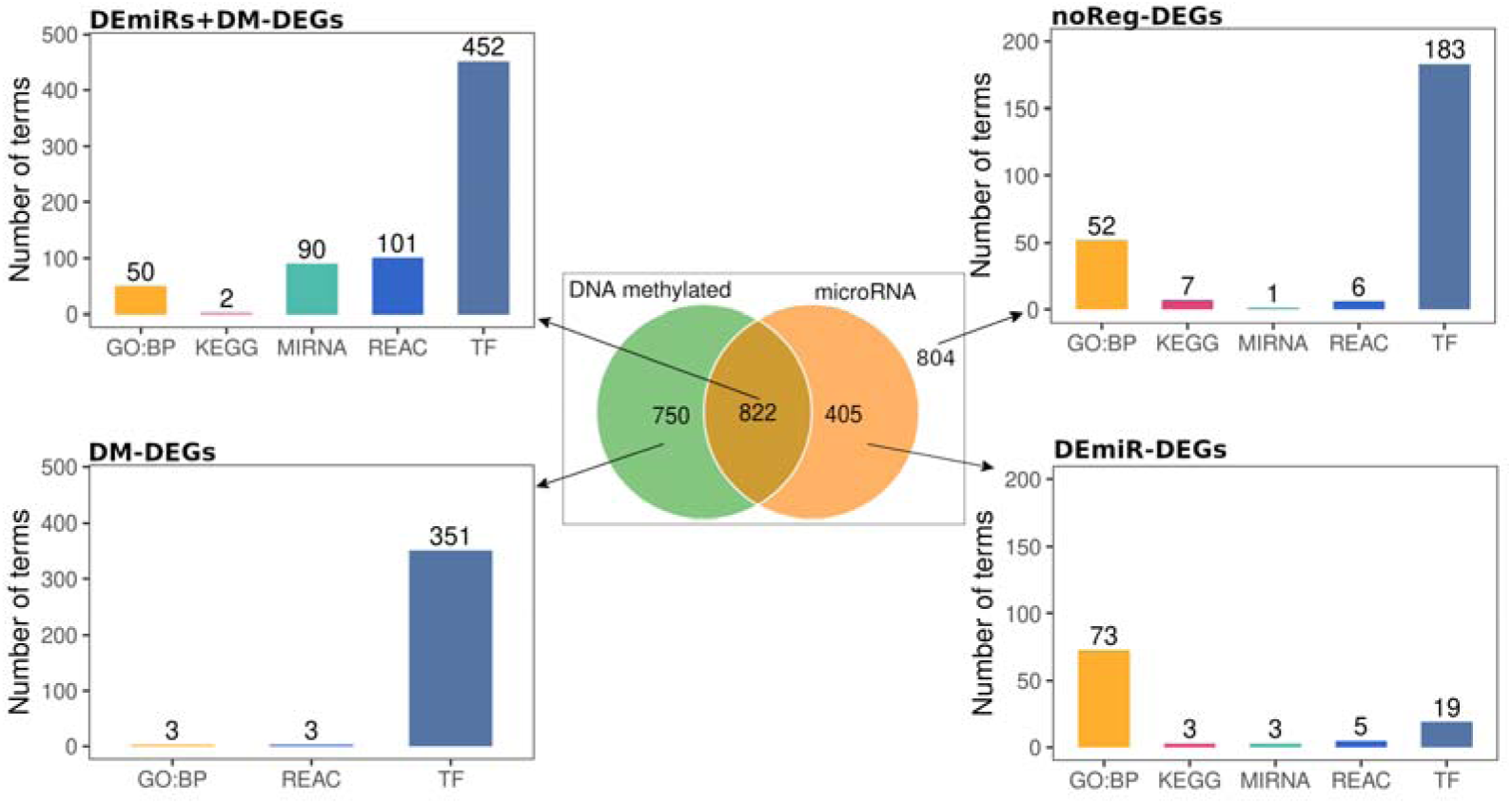
DEGs between tumour and control samples, in association with DEmiR or DM promoters. Four panels show number of terms from GSEA for each group.

Clustering of DEGs with accompanying DEmiR and DM levels revealed a critical distinction in the epigenetic factors driving HNSCC. The identified clusters predominantly demonstrated either significant DEmiRs or DM. This suggests that in HNSCC, gene expression is predominantly influenced by either miRNA regulation or methylation, but not both concurrently. This key finding illuminates the mutually exclusive roles of miRNA and methylation in shaping the gene expression landscape in HNSCC.

It is generally believed that miRNA and DNAm have an inhibitory effect on gene expression, and this pattern was evident in clusters 3, 4, 7 and 10. In addition, we observed miRNA expression and methylation patterns in clusters 1, 5, 6, and 9 that were indicative of noncanonical regulatory activity. It is important to mention that due to multiple target genes of miRNAs there was a significant overlap of underexpressed miRNAs in clusters 3 and 9 (73%) and overexpressed miRNAs in clusters 5 and 7 (77%).

GSEA analysis of each cluster linked expression and epigenetic patterns to biological function. Representative terms that best described each cluster are shown in **Figure 5** and the full GSEA results for each cluster are shown in **Supplemental table S7**. DM-DEGs in clusters 1, 4, 6 and 10 were highly enriched with genes regulated by the E2F family of TFs. Most of the genes from the E2F family were not differentially expressed in our samples (**Supplemental figure S2**), although it has been previously shown that the methylation status of CpG sites of E2F binding motif influences the binding of the transcription factor (Campanero, Armstrong, and Flemington 2000; Yin et al. 2017). In addition, cluster 6 contained underexpressed genes with hypomethylated CGIs and was enriched with genes regulated by the ZF5 transcription factor, which can function as a repressor (Numoto, Yokoro, and Koshi 1999). Clusters 4 and 5 were associated with similar terms related to various cell cycle processes but have different patterns of miRNA expression and DNAm.

**Figure 5.**
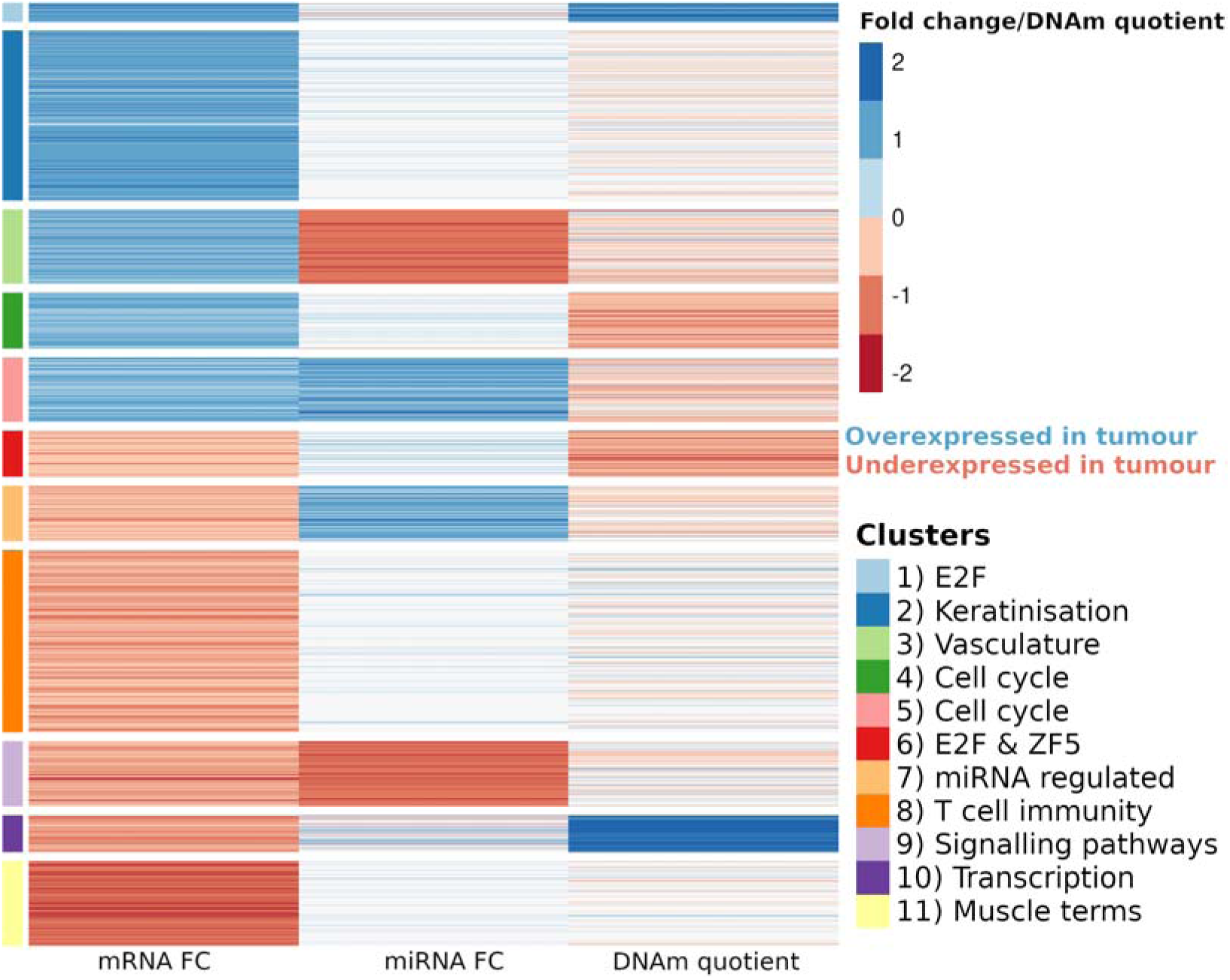
Kmeans clustering on integrated transcriptome, miRNome and methylome data on DEGs. Fold change values for transcriptome and miRNome and DNAm quotient were scaled and normalized prior to clustering. Eleven clusters are functionally described by applying GSEA over several databases (GO:BP, KEGG, REACTOME, MIRNA and TRANSFAC).

#### Influence of HPV

HPV influence on gene expression was found to be inconclusive (**Figure 2**). We observed a small number of genes and miRNAs, and no DM CGIs or sites that were significantly changed while comparing HPV positive and negative tumour sample groups (FDR-adjusted p<0.1). Number of DEGs associated with HPV status was 44, of which 25 were overexpressed (**Supplemental table S9** and **S10**). Among these, we focused on the selection of DEGs found to be functionally relevant and/or previously found to be differentially expressed in HPV- associated cancers (**Figure 6**).

**Figure 6.**
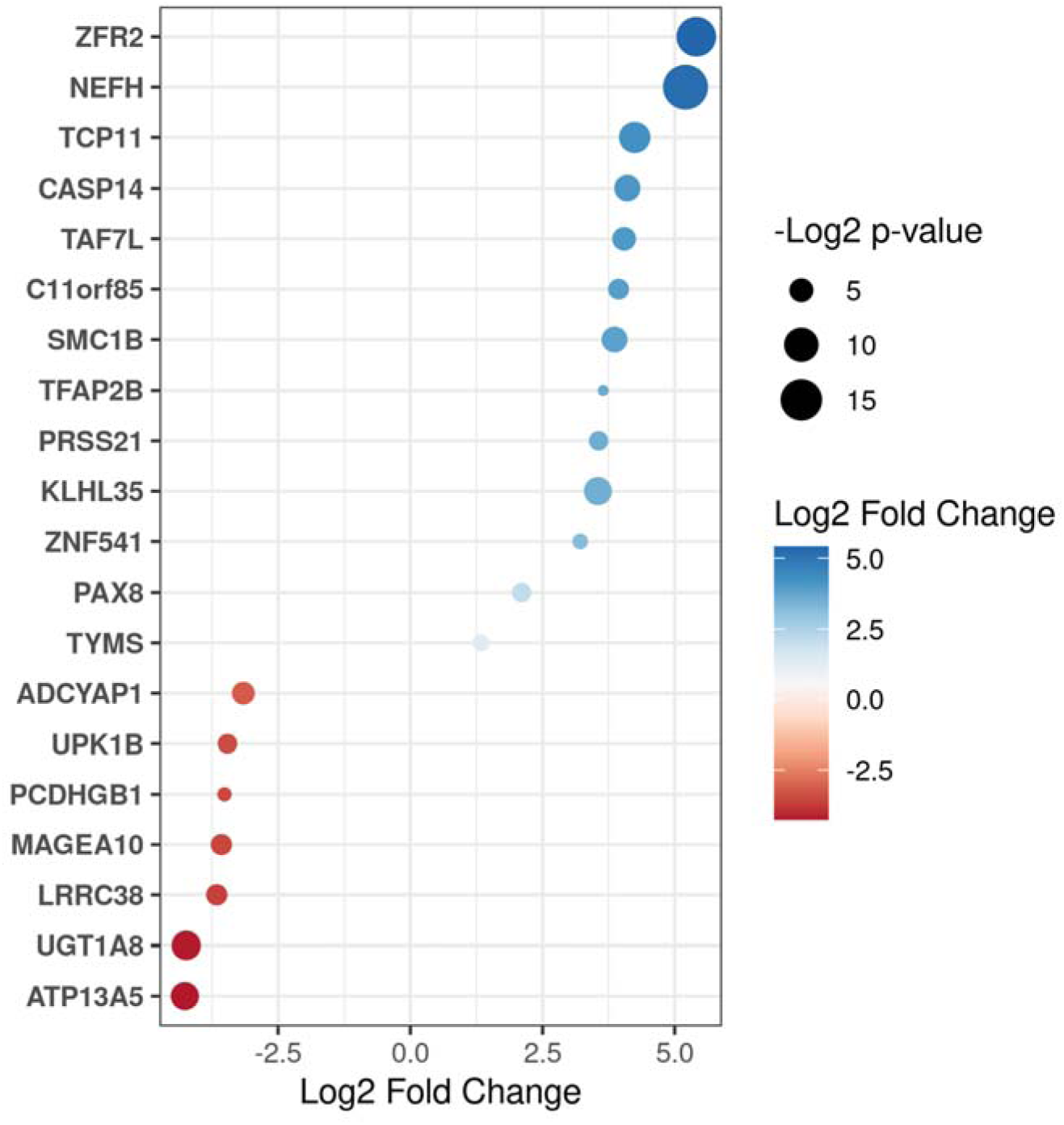
Top 20 DEGs in HPV positive compared to HPV negative tumour patients.

## Discussion

In this study, we aimed to integrate miRNome and methylome analysis with transcriptome analysis on the same samples to elucidate whether miRNAs or DNAm have more prominent impact on HNSCC development. The main goal was to identify key mechanisms and/or cellular signaling pathways of DEGs in HNSCC compared to healthy control samples.

Herein, we observed that transcriptome analysis divided samples into two major clusters: tumour and control samples, with no obvious contribution of HPV status or tumour anatomical location. Similar results were seen in the miRNA expression profile study and DNAm study in an overlapping set of samples (**Figure 2 b** and **c**) as previously published (Božinović et al. 2019; Milutin Gašperov et al. 2020). This suggests that these tumours may share similar cells of origin or common pathways during tumourigenesis.

The emergence of tumour progression pathways among DEGs (n=2781, **Figure 3**) resembles transcriptomes of radioresistant HNSCCs, where hypoxia response, p53 pathway, NF-kB pathway and inflammatory response were found abnormally activated. Interestingly, some authors found that NF-kB and other transcription factors signatures differentiate HPV- positive and HPV-negative HNSCCs.

Using unsupervised clustering, we found a cluster of genes under control of the transcription factor E2F, that were significantly underexpressed in tumour samples in comparison to controls (**Figure 5**, cluster 6). This is in line with overall hypomethylation in tumours and points to exclusive control of overexpressed miRNAs. The genome-wide hypomethylation of CpG sites is a known phenomenon that occurs in most cancers (Gama-Sosa et al. 1983; Debernardi et al. 2021). E2F family members expression was also found to be negatively regulated by miRNAs in other cancers, like lung cancers, pediatric brain tumours, and chronic myeloid leukemia. In addition, genes that were significantly associated with the E2F were found in clusters 1, 5 and 10 (**Supplemental table S6**).

In cluster 8, we found genes related to T cell immunity underexpressed in tumour samples in comparison to control samples (**Figure 5**), mostly found to be targets of overexpressed miRNAs, and in less extent associated with hypermethylation. This finding is in line with trends detected in several previous methylation studies of head and neck cancers (Moody et al. 2020; Anić et al. 2023), and in cervical cancers (Farkas et al. 2013; Milutin Gašperov et al. 2014).

Among underexpressed genes in tumours, we observed a cluster of DM-DEGs related to regulation of transcription (**Figure 5**, cluster 10). Our result showed that the cluster mostly contains genes with hypermethylation of CGIs in promoter regions and some genes that were targeted by overexpressed miRNAs. This implies that the hypermethylation of CGIs in promoter region of genes in this cluster may have disrupted binding of transcription factors required for their expression, that is a well-studied phenomena (Eden and Cedar 1994; Pikaart, Recillas-Targa, and Felsenfeld 1998; Bird 2002).

Among overexpressed genes in tumours were those that belong to cell cycle regulation (**Figure 5**, clusters 4 and 5) and vasculature (**Figure 5**, cluster 3). Herein, we found cell cycle genes have mostly hypomethylated promoter regions. On the other hand, our result showed that genes involved in vascularization in tumours are DEmiR+DM-DEGs, likely influenced by both epigenetic drivers. Moreover, other authors found a link with DNAm and cardiovascular diseases directly rather than changes driven by miRNA expression (Goossens et al. 2019), or that both epigenetic changes are involved in vascular diseases, claiming that, the most prominent gene cluster is activated via hypomethylation (Aavik et al. 2015). Interestingly, the same authors found keratin mRNAs upregulated in atherosclerosis and concluded that DNA hypomethylation may play a role in keratin gene activation and ectopic keratin expression (Aavik et al. 2015). Keratins are characteristic to epithelial cells and are not expressed in normal arteries and induction of keratin expression in atherosclerosis has been detected even earlier (Slomp et al. 1997). Our study detected similar findings in HNSCC; keratin mRNAs were found upregulated, that was partially under control of DNAm and partially under control of miRNome (**Figure 5**, cluster 2).

Herein, we found that cluster 5 (**Figure 5**) contained upregulated miRNA and their corresponding upregulated target genes. Conversely, cluster 9 displayed downregulated miRNA and corresponding downregulated target genes. This positive correlation of miRNAs and target genes has been previously observed in multiple studies (Tan et al. 2019; Xu et al. 2020). While our results could imply either a direct or indirect effect of miRNA on DEmiR- DEGs potentially through negating the action of repressive miRNA, protein complexes (Tan et al. 2019), it is important to note that our findings represent correlation, not causation. Further studies are required to establish the causal mechanisms involved. In addition, it is worth considering that miRNAs often have multiple targets, and the interactions in existing databases may be derived from high-throughput methods that may not reflect the full complexity of in vivo biological interactions.

A small number of genes were found significantly differentially expressed in HPV positive *versus* HPV negative tumours (**Figure 6**) in our study, a subset of which was previously found differentially expressed in head and neck cancer and related to HPV status, i.e. *NEFH*, *ZRF2, TAF7L, ZNF541* and *TYMS*. Notably, some authors have already found hypermethylation of *NEFH* (protein kinase binding and microtubule binding), in pharyngeal squamous cell carcinoma, which significantly correlated with HPV positivity (Nakagawa et al. 2017). *ZFR2* gene codes for RNA-binding protein characterized by its domain associated with zinc fingers and zinc ions. It was found overexpressed in HPV-positive HNSCC patients when compared to HPV-negative patients and has been flagged as prognostic for the cervical cancer cases (Tripathi et al. 2020). The expression of the transcription factor *TAF7L* (TAF7-like RNA polymerase II) has been already found increased, even 220-fold, in HPV positive HNSCCs (Slebos et al. 2006). *ZNF541* encodes a zinc finger protein supposed to be a component of chromatin remodeling complexes. The hypomethylation and higher expression of this gene was also observed in HPV-positive oropharyngeal squamous cell carcinoma (OPSCC) and significantly associated with a better overall survival, independent of HPV status (Camuzi et al. 2021). *TYMS* gene codes for thymidylate synthetase that catalyzes the methylation of deoxyuridylate to deoxythymidylate. In HPV-positive cancers they have found it overexpressed, indicating that HPV-positive oropharyngeal cancers may be more resistant to 5-fluorouracil chemotherapy than HPV-negative (Lohavanichbutr et al. 2009).

To our knowledge, we presented the most comprehensive study on an overlapping set of samples where the integration of miRNome, methylome and transcriptome analyses in HNSCC was conducted. We acknowledge certain limitations of our study, including the relatively small sample size, which may affect the validity of our findings. In addition, obtaining control samples that are genuinely representative of healthy tissue poses its own set of challenges. Associating miRNome and methylome with transcriptome profiles, we found many DEGs in HNSCC in comparison to control samples, that were mainly associated with cell cycle regulation, transcription, immunity and tumour development. We demonstrated the majority of genes belong to clusters associated exclusively with miRNome or methylome and, to a lesser extent, under control of both epigenetic mechanisms.

## Declarations

### Ethics approval and consent to participate

All patients provided informed consent to participate at the time of primary cancer treatment. The collection of samples was approved by Bioethical Bord of the Ruđer Bošković Institute (BEP-3748/2-2014) and the Ethical Board of the Clinical Hospital Dubrava (EP- KBD- 10.06.2014).

### Consent for publication

Not applicable - no personally identifiable information is presented in the manuscript.

### Availability of data and materials

Code used is available at https://github.com/PKatarina/HNSCC_Grce

Raw datasets analyzed during the current study are available in the Figshare repository under the following DOI identfiers:

Transcriptome https://doi.org/10.6084/m9.figshare.24721746

miRNome https://doi.org/10.6084/m9.figshare.24721731

Methylome https://doi.org/10.6084/m9.figshare.24721728

### All datasets and raw data will be deposited in ArrayExpress repository as well. Upon publication links will be replaced by accession numbers

Due to the file size restrictions Supplementary table S3 is deposited at Figshare repository https://doi.org/10.6084/m9.figshare.24722268

### Authors’ contributions

NMG, IS, MG and AB designed the study. KB, ED, JK, NS, IS and KM collected and processed the data. KM performed the analyses. KM, KB and NMG wrote the first draft of the manuscript. MG, IS and AB supervised and acquired funding for the study. All authors contributed to the writing of the final manuscript. All authors read and approved the final manuscript.

## Acknowledgement

The authors are thankful to Prof. Marinka Mravak-Stipetić for oral normal tissue samples collection, and to Dr. Ruth Tachezy for providing reference non-malignant tonsil control tissues. The authors are also grateful to Jasminka Golubić Talić for her technical assistance.

## Supplemental figures

**Suplemental figure 1.**
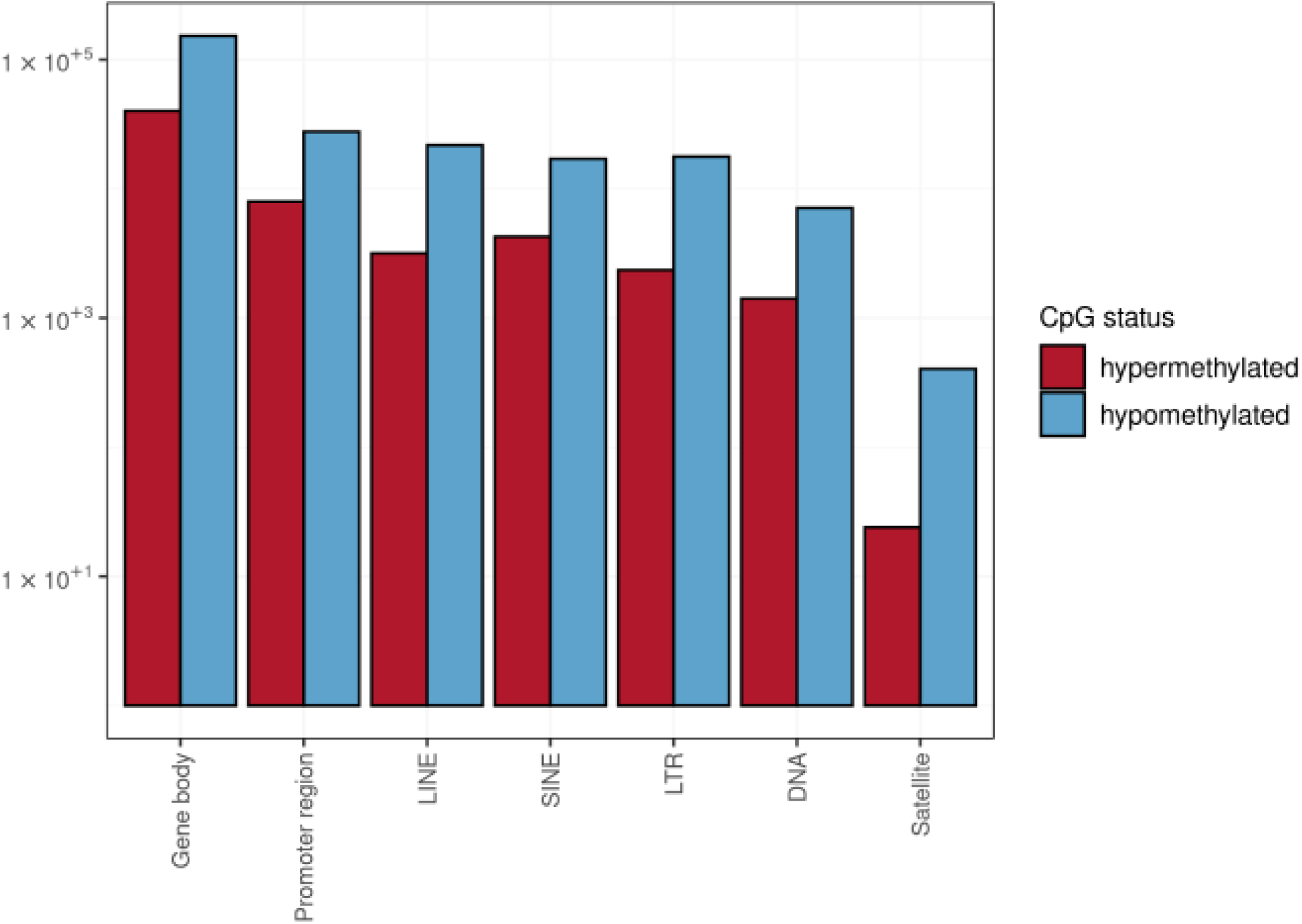
Significant number of hypermethylated and hypomethylated CpGs in cancer vs. control groups across all genomic elements (Chi squared p < 2×10^-16). Labels: Gene body – across exons and introns; Promoter region – from 500 to 2000 distance from canonical transcription start site; LINE – Long interspersed nuclear elements; SINE – Short interspersed nuclear elements; LTR – Long terminal repeat elements which include retroposons; DNA – DNA repeat elements; Satellite – Satellite repeats. Annotation for genomic elements were used from <u>Jin et al., 2015.</u>

**Supplemental figure 2.**
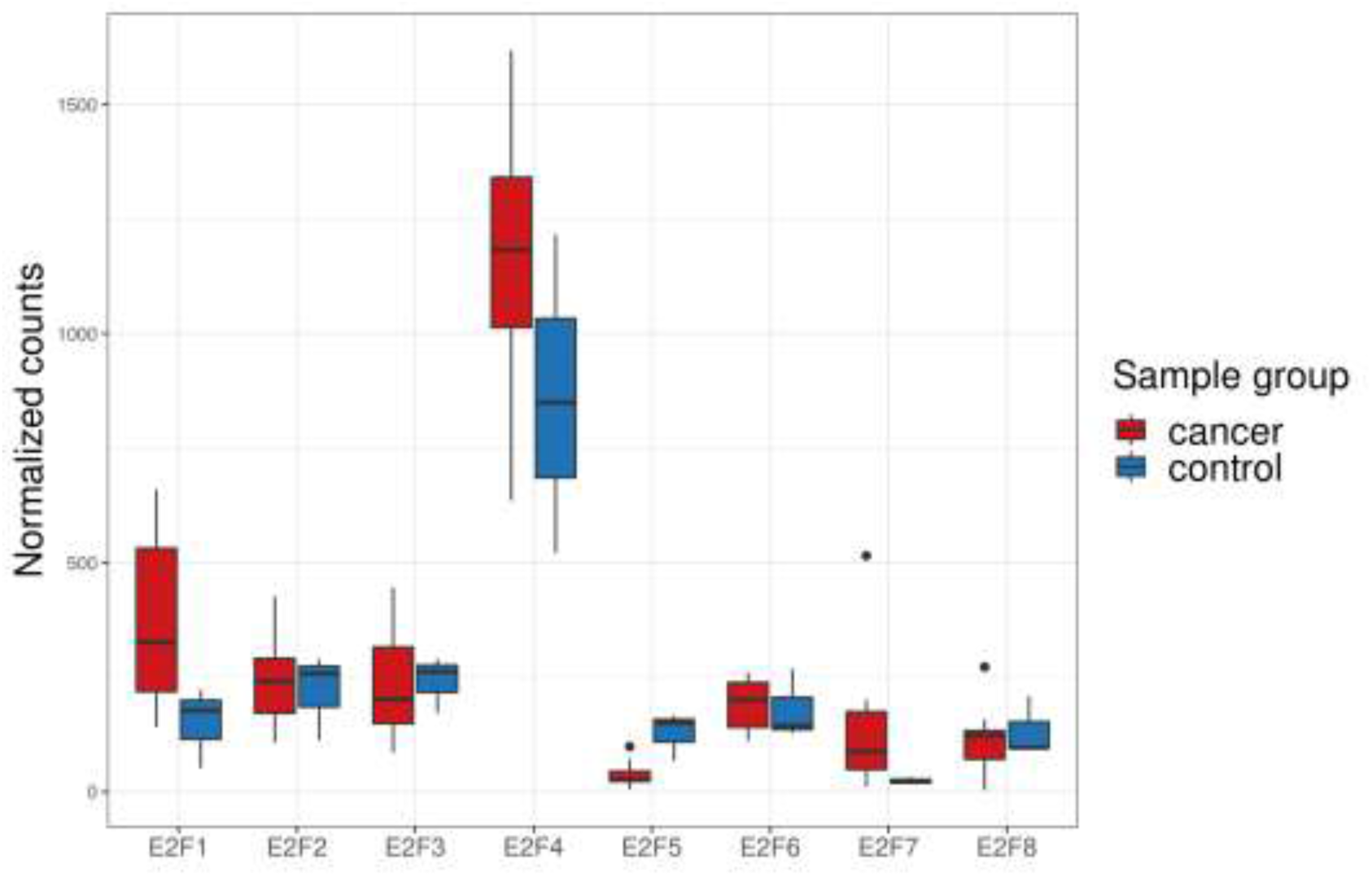
Expression of E2F family genes in cancer vs control samples.

